# Seeing around corners: Cells create chemotactic gradients to solve mazes and respond to distant cues in complex environments

**DOI:** 10.1101/836056

**Authors:** Luke Tweedy, Peter A. Thomason, Kirsty Martin, Michele Zagnoni, Laura M. Machesky, Robert H. Insall

## Abstract

Chemotaxis, in which cells steer using chemical gradients, drives fundamental biological processes like embryogenesis, metastasis and immune responses. Self-generated chemotaxis, where cells break down abundant attractants to create gradients, is an important but under-studied aspect of physiological navigation. Here we show that self-generated gradients allow cells to navigate arbitrarily complex paths and, remarkably, make accurate choices about pathways they have not yet encountered. This enables cells to solve microfluidic mazes, even with initially homogeneous environments and distant correct destinations. We combine computational models and experiments to understand how cells anticipate environmental features, and how decision accuracy is determined by path complexity, attractant diffusibility and cell speed. This permits mazes that are easy or hard for cells to resolve, despite similar appearances. Counterintuitively, slowly-diffusing attractants can generate a “mirage”, making cells prefer dead ends over correct paths. *In vivo* environments resemble complex mazes, and only self-generated gradients realistically explain cell behaviour.

## Introduction

Cells migrating in embryogenesis ^1–3^, immune responses ^4,5^ and neural pathfinding ^6,7^ steer using chemotaxis, migrating up gradients of attractants such as chemokines and netrins. Simple chemotaxis, in which gradients are established between a localised attractant source and an external sink, only provides short-range guidance ^8^. It becomes inefficient at distances above 500µm ^9^ and only functions for a narrow range of attractant concentrations. These restrictions have confounded our understanding of how chemotaxis drives longer-range phenomena such as cancer metastasis ^10^. However, when cell groups locally break down an attractant found throughout the surroundings, they create their own local, dynamic gradients ^1,11–13^, which typically direct them away from areas with a high denisty of cells, promoting metastasis ^14^. These self-generated gradients work over arbitrarily long distances, and are robust to a wide range of attractant concentrations ^8^. Here, we test their role in resolving complex paths, as for example a cell migrating through an embryo would follow. Unexpectedly, we find cells can use self-generated gradients to make accurate choices about paths they have not yet encountered. This enables cells to solve and migrate through complex mazes, even when the initial environment is homogeneous and the correct destination is arbitrarily distant. We use computational models, and live *Dictyostelium* and cancer cells in microfluidic devices ^15^, to understand how the accuracy of decisions in complex environments is determined by the complexity and lengths of the paths, and the speeds of the cells. This mechanism explains how cells can traverse longer distances than thought possible given known receptor responses, and can interpret environmental features in a way that would be impossible with simple attractant sources.

### Self-generated gradients are robust

Chemotactic cells detect attractant gradients by comparing receptor occupancy between their fronts and rears. Though cells can resolve 1% differences in occupancy ^16^, this can only guide cells for short distances – beyond 0.5-1mm, gradients are either too shallow or receptors too saturated to detect a gradient direction ^9^. One way to resolve this restriction is if the cells break down attractants while responding to them. As these dynamic gradients are usually impossible to measure directly, they are best studied using computational models verified experimentally by living cells. Fig. 1A & Movie S1 show modelled cells responding to a passive 0.5mm gradient or to a self-generated gradient in which the attractant is broken down by a cell-surface enzyme. Both mechanisms guide cells to the source in a similar time, though the self-generated gradient works faster. In contrast, cells migrate inefficiently to a 1mm passive gradient; it is either too shallow (Fig. 1C & Movie S2) or, if made steeper, rapidly becomes saturating. In comparison, the self-generated gradient gives robust chemotaxis throughout, because the gradient is always sharp and locally-produced, with a non-saturating attractant concentration maintained robustly around the cells (Fig 1C & Movie S2). Assays with real cells confirm this observation (Figs 1B & 1D, Movies S1 & S2).

**Fig 1:**
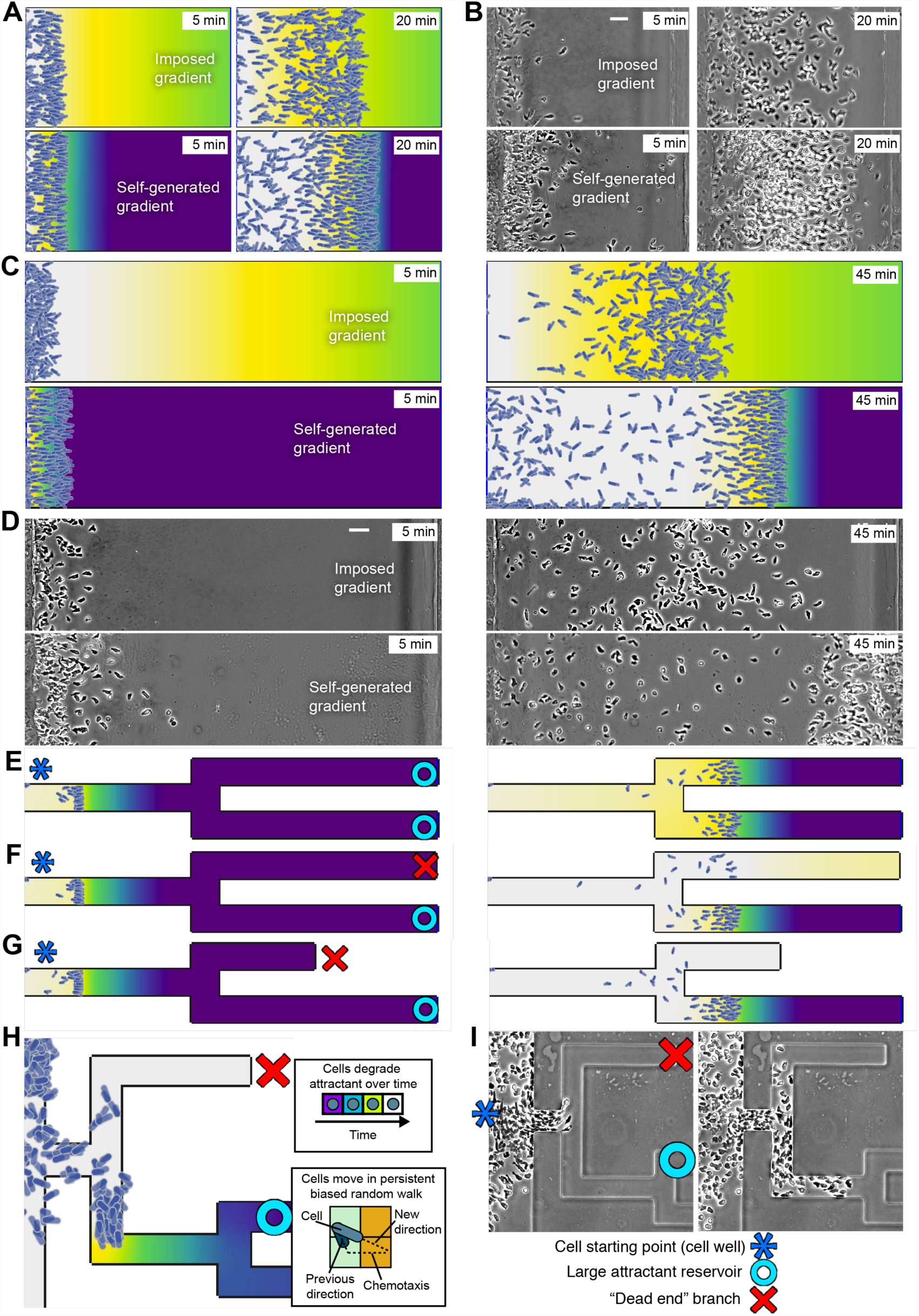
Self generated gradients allow cells to explore local environmental features remotely. (A) Comparison of imposed and self-generated gradients guiding cells across 500µm to a full attractant well. (B) Experimental verification of (A) using *Dictyostelium discoideum*. The imposed gradient uses the non-degradable attractant Sp-cAMPS. The self-generated gradient uses uniform cAMP. (C) Same as (A), but across 1mm. Here, cells following the imposed gradient perform poorly after the halfway point. In contrast, the self-generated gradient performs as well as in (A). (D) Experimental verification of (C) using the same attractants as in (B). (E-G) Simulations of a self-generated gradient navigating past a junction. In (E) There are two equal sources either side of the junction. Each branch after the junction yields the same amount of attractant, and so each recruits the same number of cells. In (F) one branch is a dead end. Some cells do still commit, as there is still a residual attractant gradient. In (G), the dead end is much shorter, and is almost entirely clear of attractant as the cells approach the junction. (H) Diagram of simulation rules. Over time, cells degrade attractant at their current location with Michaelis-Menten kinetics. Each simulated cell moves in a persistent random walk, with its new direction of motion positively biased in the direction of local attractant gradients. High concentrations can saturate a cell’s receptors. (G) Mazes are fabricated in PDMS and the behaviour of real cells is observed. This allows us to verify our mechanistic model by confirming the model’s behavioural predictions using real cells. Bar 50µm.

### Self-generated gradients allow cells to make long-range decisions

Chemotaxis studies often ignore diffusion, because they consider the steady state of an imposed, linear gradient ^17,18^ in which the diffusivity of the attractant is irrelevant. In contrast, diffusion is a key determinant of self-generated gradients. Feedback between cells depleting the attractant and migrating in the resulting gradients can yield counterintuitive results, meaning that accurate modelling and mathematical analysis of cell speed, attractant diffusivity and even local topology become essential. To that end, we used dynamic simulations to study how cells make decisions at junctions, first looking at two equivalent paths (Fig. 1E). Self-generated gradients robustly drive equal numbers of cells into each branch. Stochastic variations balance out - a branch with slightly more cells evolves a flatter gradient, directing cells into the other branch (Fig. S1A). More interesting, if one branch is closed (Fig. 1F), the cells that enter the closed branch deplete the attractant and prevent further recruitment.

When the closed branch is shorter (Fig. 1G), an unexpected behaviour emerges. The migrating cells clear the attractant from the short branch by diffusion, before they reach the junction and decide a direction. Cells thus sense the space ahead of themselves at a distance by breaking down a diffusible chemoattractant locally; they can sense a closed branch without entering it, and make accurate decisions about pathways ahead they have not visited (Fig S1B).

To understand how the cells sense closed paths without visiting them, we simulated cells following a self-generated gradient from a starting well connected by a single “correct” path to a reservoir of attractant, by analogy to physiological problems like finding a path from a tumour into a blood vessel. Parameters were taken from real measurements of *Dictyostelium* cells ^19,20^, which chemotax towards cAMP while breaking it using a cell-surface phosphodiesterase ^21^. All designs work without initial spatial cues in the chemoattractant, which starts uniformly distributed. These simulations accurately predict the behaviour of real cells in numerous experiments (figs 1,2 *et seq.*).

**Fig 2:**
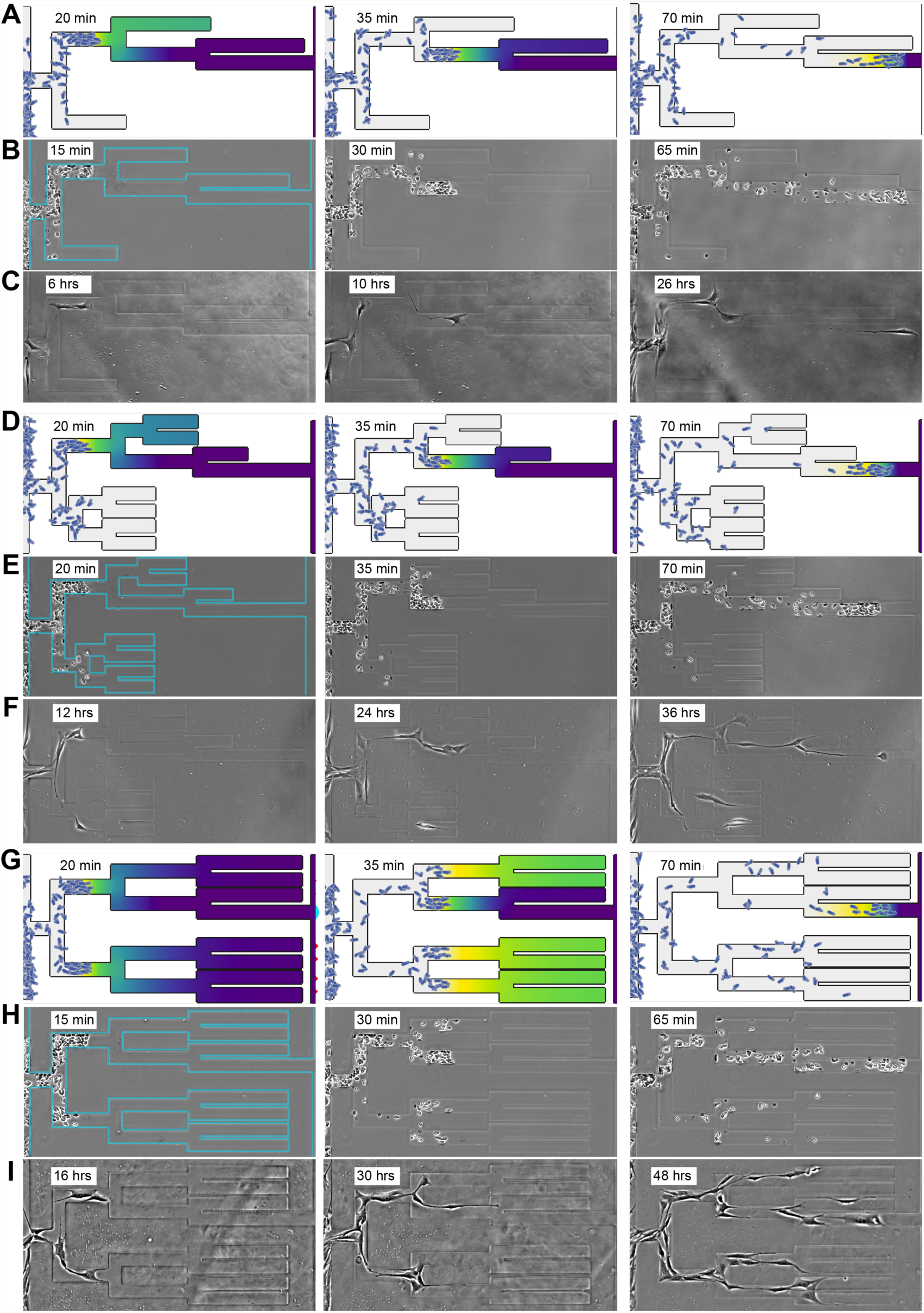
Real cells make decisions in with model predictions. (A) Simulation of cells navigating the maze with short dead ends, at time-points past the first, second and third decisions. In this design, cells are predicted to almost always commit to the correct path to lead them to the attractant well. (B) *Dictyostelium* cells migrating through the same maze design, initially filled uniformly with the attractant cAMP. (C) Pancreatic cancer cells in the same maze design, with an initial background of 10% FCS. (D-F) Simulations of the maze design in which dead ends are half the length of live ones (D), compared with experiments using the same maze design with *Dictyostelium* (E) and pancreatic cancer cells (F). (G-I) Simulations of the maze design in which dead ends are half the length of live ones (G), compared with experiments using *Dictyostelium* (H) and pancreatic cancer cells (I). Device width 500µm.

### Self-generated gradients allow cells to navigate mazes

Self-generated gradients form steeply near the cells that make them ^1,9^. This, combined with their ability to detect dead ends ahead, means they can be used to navigate environments of arbitrary length and complexity (Fig. 2). Strikingly, simulated cells not only migrate through remarkably complex structures, but also increase in accuracy as they pass through the maze, as distant branches have more time to lose attractant. Experiments with real cells in microfabricated PDMS mazes recapitulate the models with remarkable accuracy (Movies S3-S5), with the strikingly similar distributions of cells (compare Figures 2A-C, 2D-F & 2G-I). To confirm that the real gradients were self-generated, we assayed cell progress into mazes containing Sp-cAMPS, a cAMP analogue that is not broken down by extracellular phosphodiesterases ^22^. Cells made almost no progress (Fig. S1C). We conclude that self-generated gradients allow cells to accurately navigate mazes that are too long and complex for simple imposed gradients to be readable.

### Accuracy of decisions is controlled by length and complexity of paths

As the time molecules take to diffuse increases with the square of the distance, we expected the length of dead ends to control decision fidelity. Attractant in short dead ends would be depleted by the time the cells arrive at the junction. For longer dead ends, diffusion is slower, so depletion cannot establish a perceptible difference between the branches. We tested this by comparing decisions of simulated and real cells in different mazes. The model predicts that most cells will decide correctly between the path and short dead ends (Fig. 2A, Movie S3). Real cells behave the same way, with a small front pack of cells migrating through the maze with remarkable accuracy, and very few cells deviating into the dead ends (Fig 2B, Movie S4).

We measured decision accuracy in different maze designs (Fig. 2A, D & G, Fig. S2A). All had the same correct path. The first, simple maze has three short, dead-ends (Fig. 2A-C). In the second, short maze, dead ends are half the length of live ends (Fig. 2D-F). In the third, long maze, live and dead ends are nearly symmetrical, except for the final connection to the attractant reservoir (Fig. 2G-I). We focused on the first, hardest decision in each maze (Fig. 3A), recording the decision fidelity as cumulative number of cells committed to either the live or dead end (Fig. 3B). Our prediction that longer dead ends lead to poorer decision fidelity was confirmed - cells performed consistently better in the simple vs. the long maze, and better in general in the short vs. the long maze (Fig. 3C-E; blue shows cells choosing the correct path, and red the dead end). Responses for longer mazes were biphasic – initially there was little difference between the numbers of cells choosing correctly and incorrectly, but as cells depleted the attractant in the dead ends their decisions became more accurate (Fig. 3C, D). To confirm this was true for other cell types and self-generated gradients, we followed a pancreatic cancer line in homogeneous medium containing 10% FCS (Fig. 3E, Movie S5), for which lysophosphatidic acid acts as a self-generated chemoattractant ^23^; their decision-making shows the same trend as the *Dictyostelium* cells, with increasingly accurate decisions being made as the dead ends became shorter.

**Fig 3:**
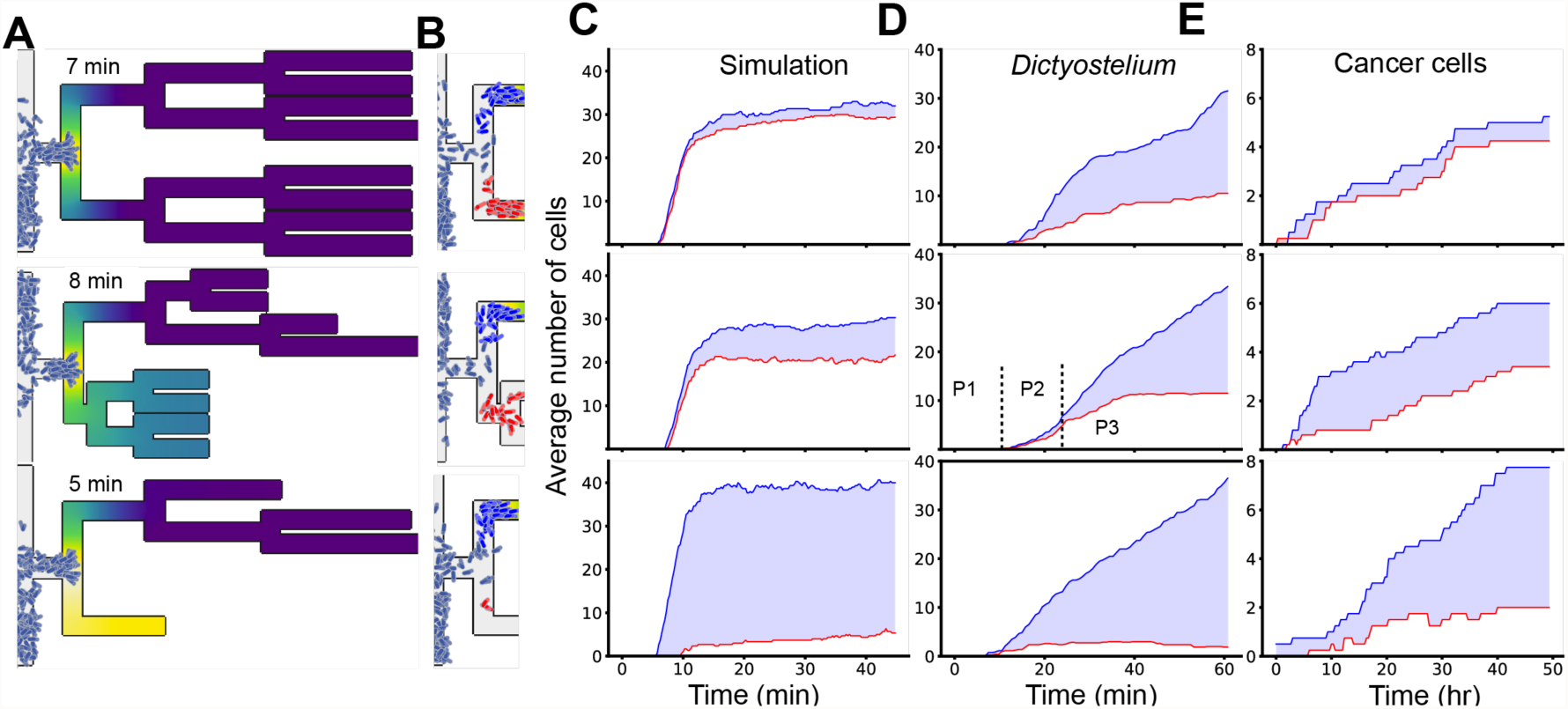
More distant and complex dead ends are harder to resolve remotely. (A) Simulations of the long, short, and pruned maze designs as the cells reach the junction between the live path and the first dead end. (B) The same simulations at a later time. Cell colour has been altered for those that have committed to the live end (deep blue) or dead end (red). Uncommitted cells are shown in the original grey blue. (C-E) Total number of cells committed to the live end (blue) and dead end (red) over time in three repeats of the same simulation environment (C), *Dictyostelium* experiments (D) and pancreatic cancer experiments (E), with t=0 when the cells first reach the entrance to the maze. Light blue shading shows the difference between these values, with a thicker light blue area showing that more cells have committed to the live path (connected to the attractant well). (E) Pancreatic cancer cells moving through a maze in response to 10% fresh FCS.

### Cell speed and attractant diffusivity

We supplemented our modelling and experimental approaches with a mathematical analysis of a decision at a T junction, with variable live and dead end length and time taken before making a decision. As real cells are not static when they decide direction at junctions, we employed a mathematical trick to map cell speed onto waiting time and verified it against our simulations (see SI section 3.1). This yielded three key predictions (Fig. 4A):

1. Decisions are better for shorter dead ends,
2. Decisions are better for shorter live ends,
3. Decisions are better if cells take longer to make them.

**Fig 4:**
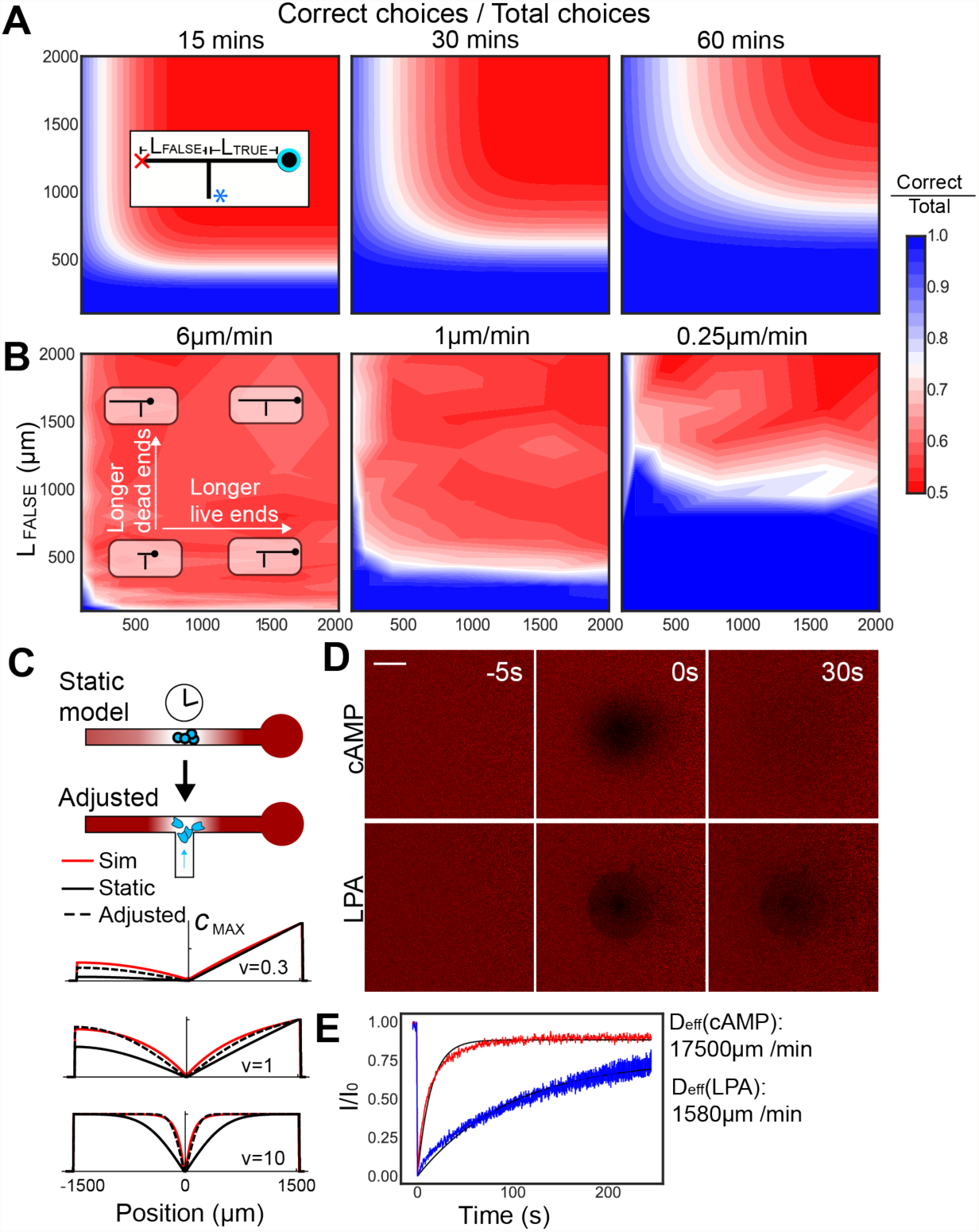
Slower cells and faster diffusion lead to better decision fidelity (A) Mathematical model of cell decisions at a T junction. Each panel shows a snapshot after a different waiting time. The x and y axes show the lengths of the live and dead ends respectively. (B) Decisions made by simulated cells moving at different speeds, and therefore taking less time (leftward) or more time (rightward) to reach the junction. (C) To correct for cell movement sharpening the gradient, we create a mapping from the solvable static model to an adjusted model. The static and adjusted models are shown for three cell speeds against the true gradient seen in the simulations. (D) Photobleaching of 8 fluo-cAMP and TopFluor lyso-PA. (E) Recovery curves & fits for the photobleached areas suggest that the effective diffusivity of LPA is ~1/10^th^ that of cAMP. Bar 50µm.

We had already observed the first prediction experimentally, strengthening our trust in the others. The second happens both because equilibrium is quickly reached, and because the resulting well-to-junction gradient is steeper. The third is due to cells having longer to clear attractant diffusing out of dead ends prior to deciding which path to take. This explains the greater efficiency of the second and third decisions in the mazes in movies S3-5. The pattern is observed when altering cell speed in simulations, as slower cells effectively have longer to make decisions (Fig. 4B; note similarity to 4A).

These findings raise an apparent contradiction. We predict that slower cells will make better decisions than faster ones, yet *Dictyostelium*, which solve mazes in two hours, perform similarly to cancer cells that take roughly two days. As their respective attractants (cAMP ^24^ and LPA ^14^), have similar molecular masses, we had expected them to have similar diffusivities. However, lipids are often carried by proteins such as albumins, which slow their effective diffusion. We predicted the relationship between the diffusivities of cAMP and LPA that would lead to equal decision fidelity, and then performed a photobleaching assay on fluorescently labelled LPA and cAMP (Fig. 4E,F). The effective diffusivity of LPA was strongly reduced (to about 1/10^th^), as expected, with the relationship between the molecules similar to that predicted mathematically (about 1/20^th^, SI Section 3.2).

### Complex topologies drive cells into incorrect decisions

The behaviour of real cells raised one other mystery. Our analyses show dead end length is crucial to decision fidelity. Despite this, cells in the short maze did not outperform those in the long maze until late in the experiment (movie S4). In fact, between the cells approaching the decision point (P1, Fig. 3D) and correctly committing to the live end (P3), there seems to be a short period where the decisions are worse in the short than in the long maze (P2). To understand this, we modelled attractants with a range of diffusivities, expecting that high diffusivity would yield excellent decisions, and that fidelity would monotonically decrease until decisions were made with no information and 50% of cells head in each direction. Decisions were accurate at high diffusivities (Fig. 5A, Movie S6), decreasing to near 50:50 at normal small molecule diffusivities and for extremely low diffusivities (Fig 5B), but unexpectedly, at diffusivities around 1/10th that expected of a small molecule like LPA or cAMP, we found that decisions were predicted to be worse than 50:50 for a time, with a majority of cells committing to the dead end (Fig. 5B). Analysis showed that the counterintuitive result derived from the complexity of the path. Two or more branches from the dead side of the maze were supplying attractant molecules, whereas in the live end molecules travelling a similar distance only came from a single branch. More distant upstream bifurcations had a minimal effect. This leads to a surprising conclusion - complex, branched environments can create short-term chemotactic mirages, which lead cells away from the source of attractant (Fig. S4).

**Fig 5:**
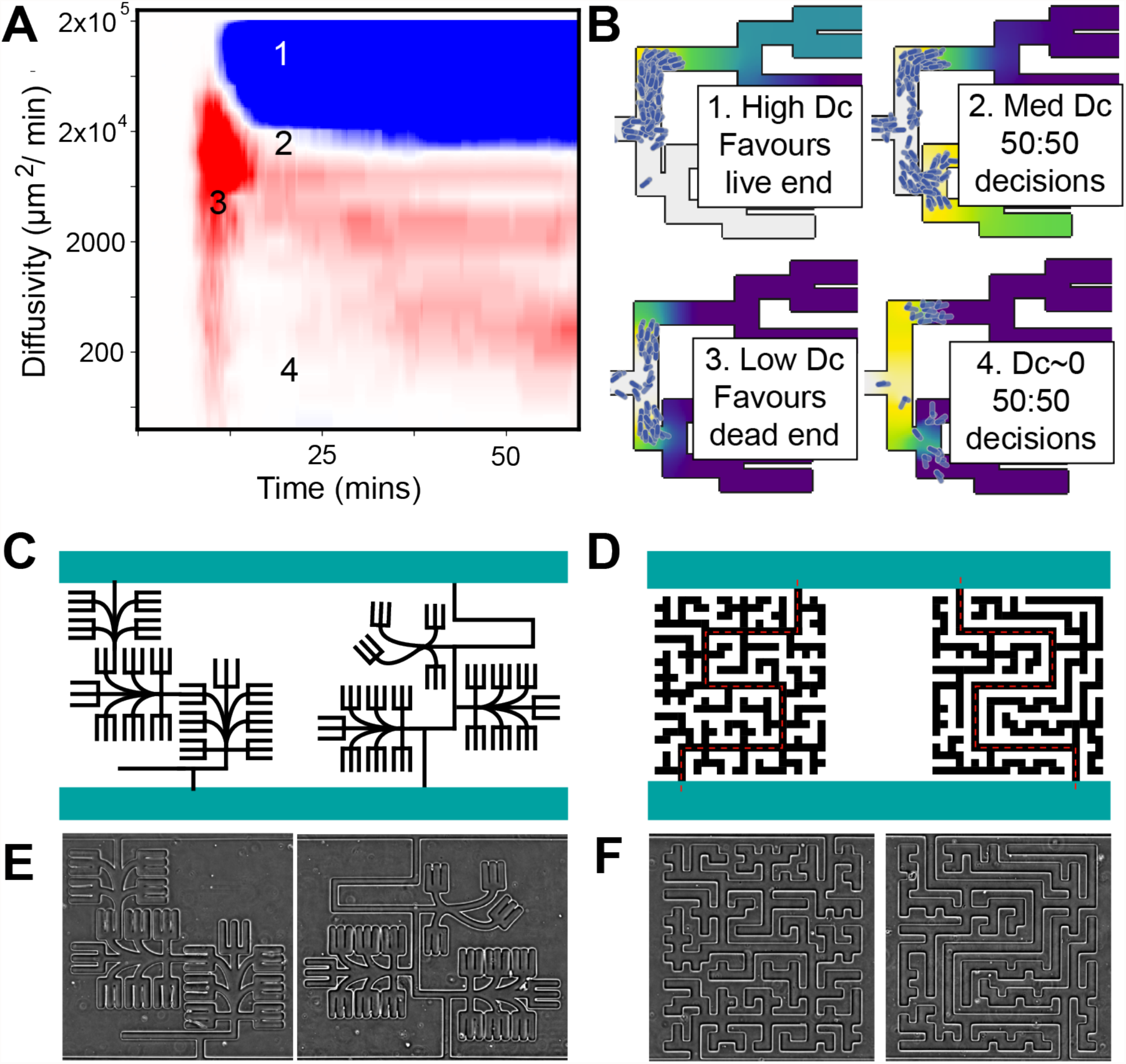
Cell decisions are a function of the attractant amount transported to them. (A) Decision fidelity over time for the short maze, simulated using various diffusivities. Four areas of note are labelled, showing phase changes in decision behaviour. (B) Still frames of the four labelled points in (A). Of particular note is 3. in which more cells commit to the dead end than to the live end. (C) Designs for easy (left) and hard (right) trident mazes. Though both have the same amount of dead end space, the hard maze is designed with fewer, longer, and more branched dead ends. We note that the easier maze is harder to solve visually. (D) Designs for easy and hard labyrinths, along similar principles to (C). The solution is marked with a red, dotted line. (E, F) Phase contrast images of the four maze designs after fabrication. Device height (top to bottom in panel E) 500µm.

To test whether chemotactic mirages affect real cells, we fabricated pairs of closely-related maze designs (Fig. 5C & D) with comparable paths to the source, but with one “easy” (with models predicting accurate decisions) and one “hard” (Movies S7-8; Fig. 6A-D). Cells responded extremely divergently, as predicted, despite the mazes’ superficial similarity. They overwhelmingly chose accurately in the “easy” maze (Fig. 6A), with simulations showing low levels of attractant in the first dead end as cells arrive. In contrast, cells in the corresponding “hard” maze design (Fig. 6B) showed an extended period of 50-50 decision making followed by a weak commitment to the correct path. In simulations, this is due to the similarity of the attractant gradient leading up the live and dead ends; when the dead end starts to clear, the cells regain accuracy. The second maze pair was even more striking. Cells overwhelmingly chose correctly in the “easy” maze (Fig. 6C), due to early attractant depletion down the dead end. In the “hard” maze, there was a time window in which cells committed more to the dead end than the live end (Fig. 6D, area shown in red), demonstrating that the chemotactic mirages predicted by models indeed affect live cells. In each case, a substantially greater fraction of the cells choose inaccurate paths in “hard” than “easy” mazes (Fig. 6E). Mathematical analysis shows that when distances to attractant sources are longer (3mm or more), mirages could dominate (Fig. S5).

**Fig 6:**
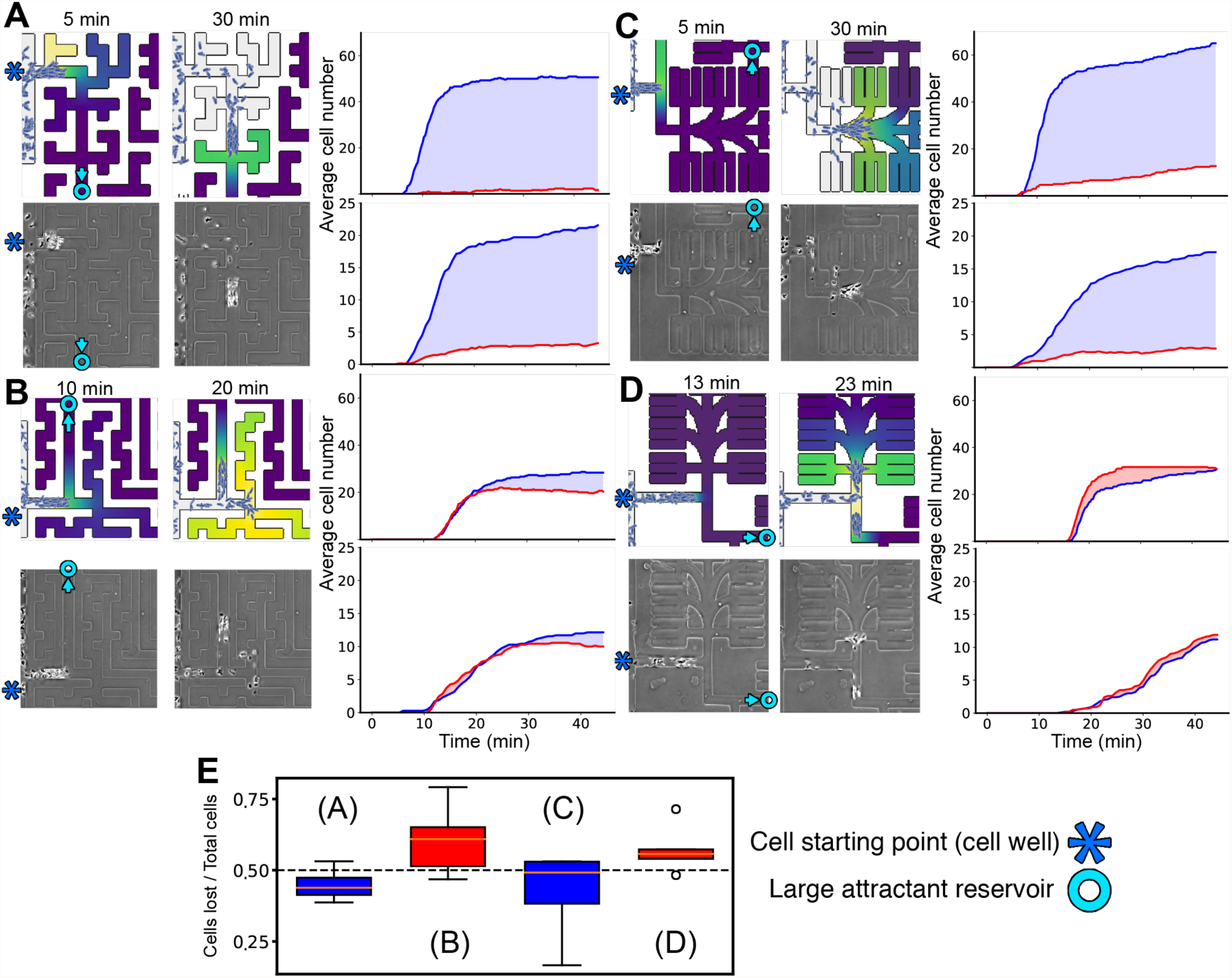
Deliberate misdirection of live cells by designing the environmental topology. (A) Decision results for the easy labyrinth. Stills at, and after the first decision show similar behaviour between the real and simulated conditions, with overwhelming commitment to the live path. The simulation still at the decision point shows that this is due to the weak residual attractant gradient in the dead end. (B) Decision results for the hard labyrinth. The period of poor decision making is prolonged in this case. The simulation stills show that the dead end provides a steep gradient as the decision is made, but that this quickly disappears after cells pass the junction. (C) Decision results for the easy trident maze show live end commitment, as with (A). (D) Decision results for the hard trident maze. In this case, both in the simulation and the experiments, more cells commit to the dead end than to the live one. (E) Final positions of cells following the final decision in the hard mazes (or, having travelled a similar distance in the easy mazes). Cells are more likely to have become committed to (and lost in) a dead end in both hard maze designs than their easy counterparts (n=4, whiskers show 1.5x interquartile range; points outside shown singly). All devices 500µm wide.

## Discussion

This work has clear and wide-ranging implications for biology in general. We find that the details of cells’ environments strikingly affect their ability to steer accurately, and allow them to steer using information it seems impossible for them to have obtained. Changes that seem minor make a substantial – and predictable - difference to the accuracy and eventual destination of the cells. By matching our experimental results to mathematical and computational methods, we ensure that our findings are general, applying to any system employing an attractant that can be degraded by the cells that respond. This mechanism differs entirely from that used by *Physarum polycephalum* to solve mazes ^25^, which relies on the plasmodium migrating down all branches simultaneously before pruning useless paths ^26^.

Many situations where chemotaxis occurs *in vivo* – neutrophils extravasating to an infection in tissue ^27^, for example, or germ cells migrating through an embryo ^28^ – have equivalently complex paths. Similarly, attractant degradation is almost universal, with examples known in immunity ^29^, development ^28^ and cancer ^30^. Chemotactic cells have a variety of well-known mechanisms for depleting attractants, including receptor-ligand endocytosis ^31^, decoy receptors ^1,11,32^, and cell-surface enzymes that degrade attractants ^30,33^. Ligand breakdown is rarely considered when interpreting spatial patterns of chemotaxis data - its effects can be complex, counterintuitive and difficult to measure - but it needs to be. A combination of computational modelling, as shown here, and modern microscopical techniques is required.

We have described decisions as better or worse, but it is crucial to note that these loaded terms contain a value that does not apply to an *in vivo* context. Our aim is to understand how complex topology might draw them to places that intuition would not predict. The chemotactic mirage, in which strongly branching topologies can demonstrate more powerful chemoattraction than an actual source of chemoattractant, is particularly counter-intuitive. This could be crucial to understanding metastatic behaviour in complex *in vivo* environments ^10^; the metastasis of glioblastoma, for example, is known to move along the long, highly branched white matter tracts of the brain ^34^. Overall, the detailed interactions between cells’ breakdown of chemoattractants and the geometry of the paths they chemotax along are fundamentally physiologically important, and need to be studied with the same intensity as attractant sources.

## Methods

### Maze design and fabrication

Design schematics for mazes are given in the SI. Microfluidic mazes were fabricated in polydimethylsiloxane (PDMS) (Sylgard 184, Dow Corning, US) using standard soft lithography techniques. Briefly, silicon masters were produced using SU8 3005 photoresist (3000 series, MicroChem, US) on silicon wafers following the manufacturer’s protocol to achieve a final resist thickness of either 10 or 15 µm. The resist was exposed through a photomask (JD Photo-Tools, UK) to collimated UV light and was developed in MicroPosit EC solvent (Rohm and Haas, US). To prevent PDMS adhesion to the silicon master, this was salinized by vapour deposition of 1H,1H,2H,2Hperfluorooctyl-trichlorosilane for 1 hour. PDMS was poured onto the silicon master at a 10:1 ratio of base to curing agent, degassed in a vacuum desiccator chamber and cured at 70°C for at least 3 hours. PDMS devices were then peeled from the mould, cut to the desired size and 2mm holes were punched to obtain inlet and outlet ports. PDMS devices were then cleaned and irreversibly bonded to glass-bottom petri dishes (manufacturer) using oxygen plasma.

### Maze use

Mazes were filled uniformly with medium by filling all inlet ports with the medium of choice (typically ~6µl per well) and then placing into a vacuum desiccator for around 20min, degassing the PDMS. When the vacuum is released, the pressure difference draws medium into all parts of the maze, including dead ends, although this functions best if additional medium is pipetted up and down in each well, as this dislodges residual gas bubbles. Mazes used for cancer cells were pre-filled with 0.05% BSA in sterile, deionised water in order to block the PDMS and prevent any attractants from adhering. The pre-fill was dried out by first draining the wells thoroughly with a pipette, then placing in a tissue culture hood for ~2h. As soon as a maze was observed to be dry, it was re-filled with experimental medium (which, for cancer cells was Dulbecco’s Modified Eagle Medium (DMEM)+10% FCS, freshly added).

### Cell Lines

*D. discoideum* wild-type line NC4 were used for experiments in the long, short and simple mazes. Cells were starved on agar until scrolling waves were observed (typically taking 4-5hrs), then harvested in the maze medium (Development Buffer (DB) + 1µM cAMP + 3mM caffeine), and inoculated into the cell well at high density. An adenyl-cyclase null (*aca-*) in an AX2 background was used for experiments in the easy vs hard mazes. Cells were developed by 1.5hrs starvation in DB, followed by pulsing with 300nM cAMP every 6 minutes for 4.5hrs. In both cases, the *D. discoideum* were grown non-axenically on bacterial lawns. The cancer line used was KPC model murine pancreatic cancer (-kras -p53). Cells were cultured in DMEM+20%FCS, then trypsinised, resuspended in DMEM+10% fresh FCS and placed in the cell well of the maze. In these experiments, mazes were filled with DMEM+10% fresh FCS.

### Simulations

Simulations were written in Java. Diffusion in a complex environment was simulated using the semi-implicit DuFort-Frankel method. Agent-based model cells then made decisions using a persistent, biased random walk.

## Data availability

**No data sets were generated or analysed during the current study.**

## Code availability

**The source code will be made permanently available on GitHub upon publication**.

## Acknowledgements

We are very grateful to Dr. Heather Spence for assistance with tissue culture. Funded by a core grant (A24450) and a multidisciplinary project award (A20017) from Cancer Research UK.

